# Gene networks provide a high-resolution view of bacteriophage ecology

**DOI:** 10.1101/148668

**Authors:** Jason W. Shapiro, Catherine Putonti

## Abstract

Bacteriophages are the most abundant and diverse biological entities on the planet, and new phage genomes are being discovered at a rapid pace from metagenomes. As more novel, uncultured phage genomes are published, new tools are needed for placing these genomes in an ecological and evolutionary context. Phages are difficult to study with phylogenetic methods, because they exchange genes regularly, and no single gene is conserved across all phages. Instead, genome-level networks have been used to group similar viruses into clusters for taxonomy. Here, we show that gene-level networks provide a high-resolution view of phage genetic diversity and offer a novel perspective on virus ecology. To that end, we developed a method that identifies informative associations between a phage’s annotated host and clusters of genes in the network. Given these associations, we were able to predict a phage’s host with 86% accuracy at the genus level, while also identifying genes that underlie these virus-host interactions. This approach, thus, provides one of the most accurate means of host prediction while also pointing to directions for future empirical work.

## Introduction

Bacteriophages (phages) are viruses that infect bacteria, and with over 10^31^ estimated on the planet, are often the most abundant and diverse members of any ecosystem (Edwards & Rohwer 2005). Phages act as predators, drivers of biogeochemical cycles (Wilhelm & Suttle 1999), industrial contaminants (McGrath et al. 2007), and as important mutualists within bacterial pathogens that cause disease in plants and animals (e.g. Addy et al. 2012, Waldor & Mekalanos 1996). Phages have also been used as therapeutics in agriculture (Greer 2005) and for treating antibiotic-resistant bacterial infections (Chan et al. 2016, Biswas et al 2002). Similar to bacteria, the majority of phages cannot be propagated in the lab, either because their host cannot be grown or because their host is not known. Nonetheless, new metagenomes from diverse environments are being published regularly, and the rate of uncultured, novel virus discovery has increased rapidly in the past decade (e.g., Simmonds et al. 2017, Paez-Espino et al 2016, Bruder et al. 2016, Roux et al. 2015, Dutilh et al 2014). Coping with this deluge of data requires new computational methods for both classifying virus diversity and for inferring key features of virus ecology and evolution.

Except for strain-level variation of a particular virus, traditional phylogenetic methods cannot be applied to derive a “species” tree for phages. There are no universal genes shared by all phages, and horizontal gene transfer (HGT) between viruses is common. In essence, every phage genome is a mosaic that reflects the often disparate evolutionary histories of its genes (Pedulla et al. 2003, Hendrix et al. 1999), and genome-level classification is, therefore, difficult. To overcome these challenges, network-based approaches have been used to depict the relationship between phage genomes on the basis of the similarity of their genic content or overall sequence identity (Cresawn et al. 2011, Halary et al. 2010, Lima-Mendez et al. 2008, Roux et al. 2015, Paez-Espino et al. 2016).

Genome-level network analyses are appealing, because they make it possible to visualize phage relationships in place of traditional phylogenies (e.g. Paez-Espino et al 2016, Lima-Mendez et al 2008). At the same time, these whole-genome analyses continue to ignore the mosaic architecture of phage genomes and take the focus away from the actual targets of selection: genes. As a result, it is unclear how to apply these genome networks to questions beyond taxonomy. In the present work, we instead build a network of genes, where genes are connected if they are ever found within the same genome. By extending network analyses from genomes to genes, it is possible to address questions directly related to virus ecology and evolution, such as how particular genes affect the mode of infection, virulence, and host range of a virus.

Host range, in particular, constrains viral ecology and evolution, and predicting a virus’ host is a key challenge when characterizing novel, uncultured genomes. Host range typically depends on individual virus-host gene interactions (Labrie et al. 2010), and both phages and their hosts can acquire genes that alter these interactions through HGT (Meyer et al. 2016, Sachs & Bull 2005, Tzipilevich et al. 2016). Methods for predicting virus host range from genomes commonly rely on comparing genomic properties such as k-mer frequencies, codon usage, or, when possible, host CRISPR content. The best of these methods, however, are rarely better than 80% accurate for predicting a phage’s host at the genus level (Ahlgren et al. 2016, Villaroel et al. 2016, Edwards et al. 2016). Here, we build a gene-level network representing the co-occurrence of genes across phage genomes. In addition to providing a robust view of virus genetic diversity, clusters within this network can be associated with virus host range. By identifying genes that increase the correspondence between phages and their hosts, we are able to predict virus host range at the genus level with over 85% accuracy for many host genera.

## Building Genome- and Gene-Level Networks

We built genome- and gene-level networks for a set of 945 phage RefSeq genomes, consisting of 92,801 gene sequences. In the genome network (Figure 1a), nodes represent virus genomes, and two nodes are connected if they share at least one gene. In the gene network (Figure 1b), nodes represent homologous phage protein sequences, and two nodes are connected if these genes are found in the same genome. Homologous genes were identified with as low as 35% identity via clustering by usearch (Edgar 2010). Singleton and doubleton clusters were removed from consideration to increase the reliability of connections between genes. This filter yielded a final set of 8,847 gene clusters from across 913 phage genomes, dropping 32 phage genomes from primarily under-sampled, tailless phage families, which are often underrepresented in metaviromes (Steward et al. 2013).

**Figure 1.**
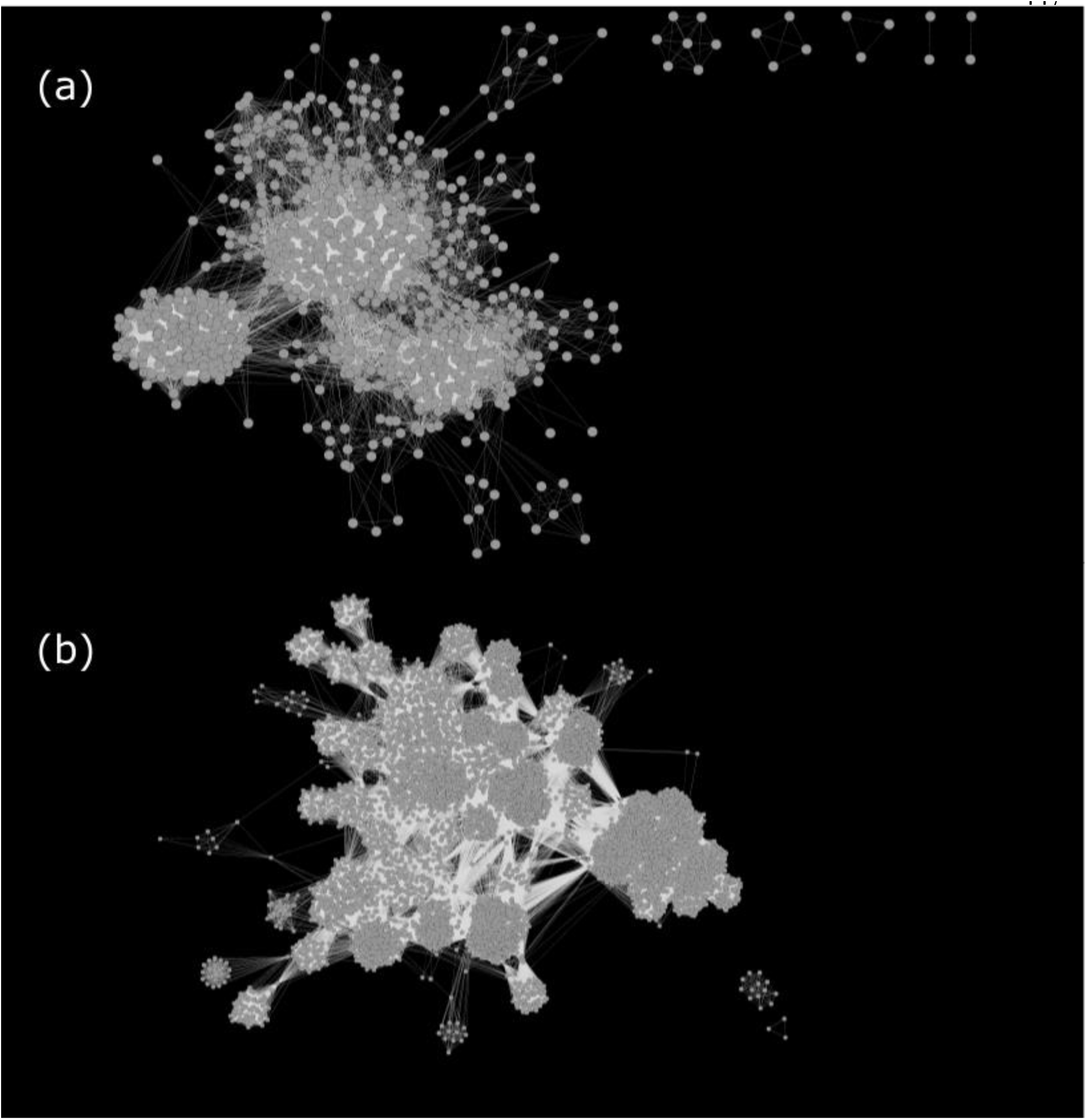
Genome-level (a) and gene-level (b) networks for a set of 913 phage. In the genome network, nodes are genomes, and two nodes are connected by an edge if they share any genes. Inversely, in the gene network, nodes are genes, and two nodes are connected if they are found in the same genome.

In each network, there exist subsets of nodes that form subgraphs in which members have more connections in common with each other than with the rest of the network. We formally identified these subsets of interconnected nodes using the Markov Clustering Algorithm (MCL) (Enright et al. 2002). MCL relies on an inflation parameter that transforms the adjacency matrix of the underlying network. Higher inflation values generally yield more clusters from a network, and others have previously used a measure of cohesion within subgraphs, the “intracluster clustering coefficient” (ICCC), to optimize this parameter choice for virus taxonomy (Roux et al. 2015, Lima-Mendez et al. 2008). Using this metric, we chose an inflation factor of 6 for the genome network and 4.1 for the gene network (see Figure S1). These values correspond to 209 clusters in the genome network and 135 clusters in the gene network. As seen in Figure 2, the MCL clusters in the gene network appear to provide a cleaner visualization of virus diversity than clusters in the genome network.

**Figure 2.**
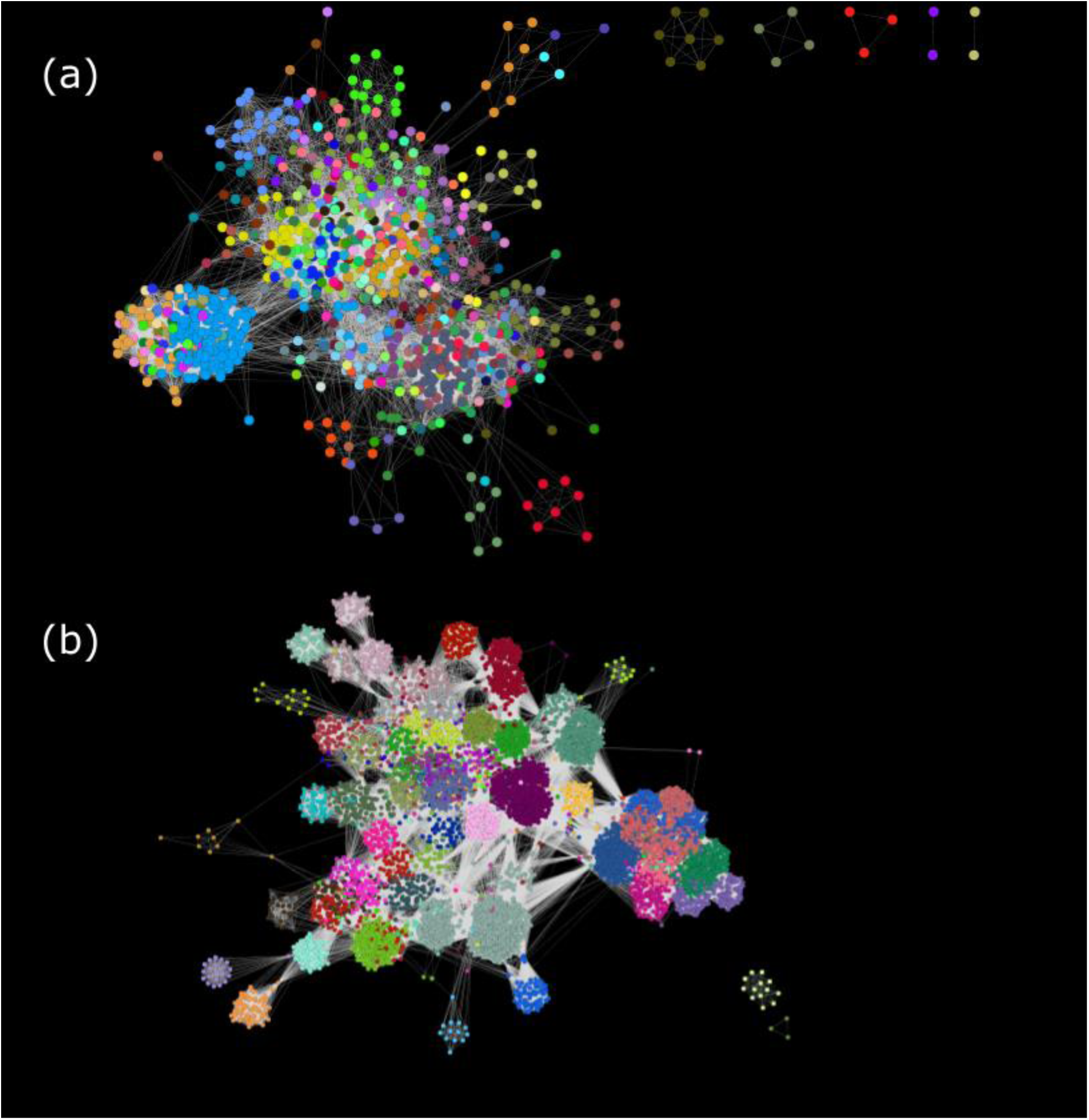
The genome (a) and gene (b) networks are identical to those in Figure 1, except nodes have been colored based on their membership in graphical clusters identified using MCL with inflation set to 6 for the genome network and to 4.1 for the gene network.

## Clusters of phage genes are associated with phage host genera

Given the gene and genome networks, we then recolored the nodes according to the phage host genus (Figure 3). In the gene network, each node represents a set of homologous genes, and only the most common host associated with these homologs is indicated for each node. As can be seen in Figure 3, phage host was poorly associated with graphical clustering in the genome network but maps closely to graphical clusters in the gene network. In fact, for several hosts, distinct clusters could be identified in the gene network that correspond at the species or strain-level of the phage host (see Figure 4).

**Figure 3.**
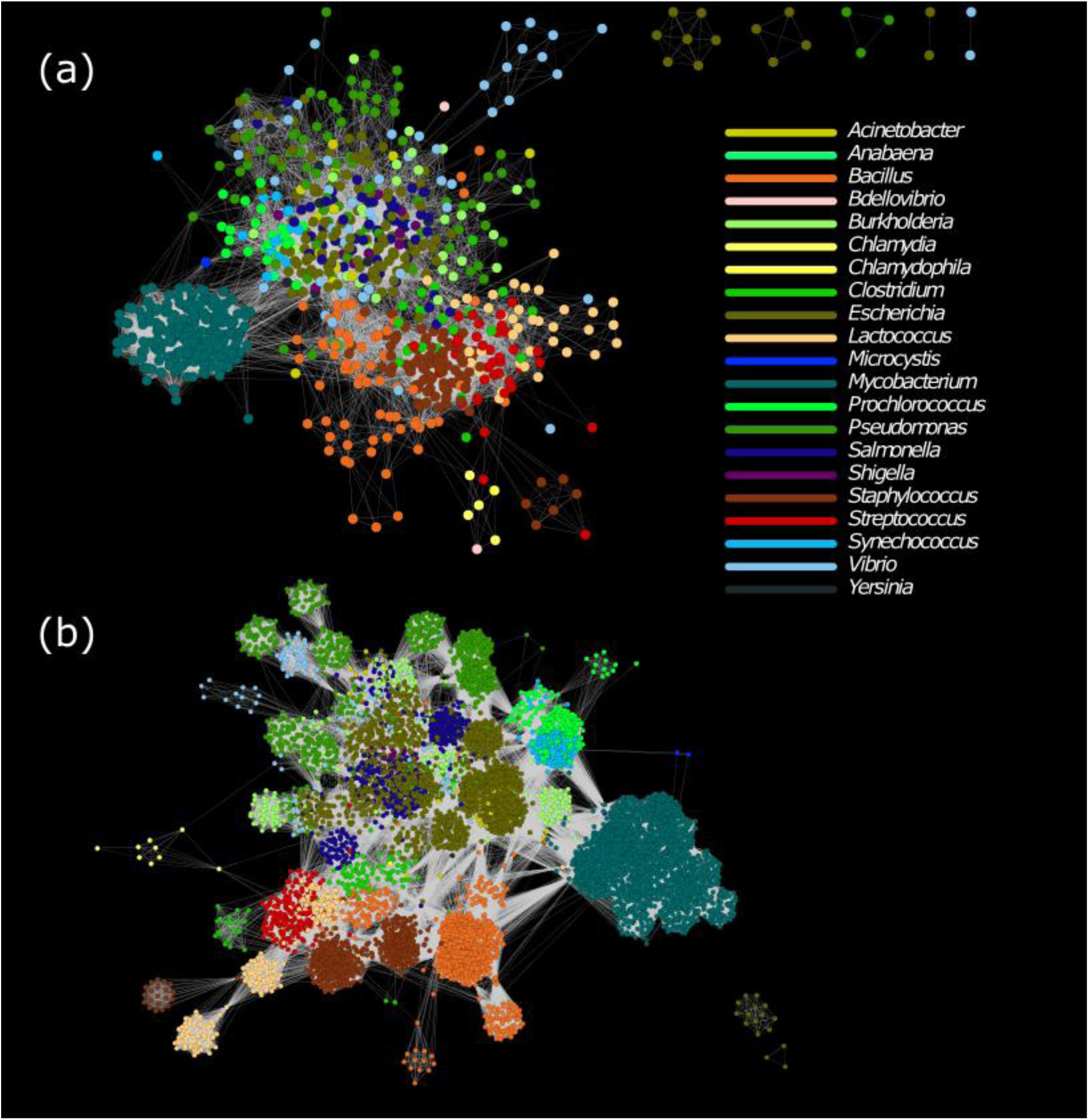
The genome (a) and gene (b) networks are identical to those in Figures 1 and 2, except nodes have now been colored to reflect the host genus associated with each phage. In the gene network, each node signifies a set of homologous sequences, and colors match the most common host for the genomes containing these homologs.

**Figure 4.**
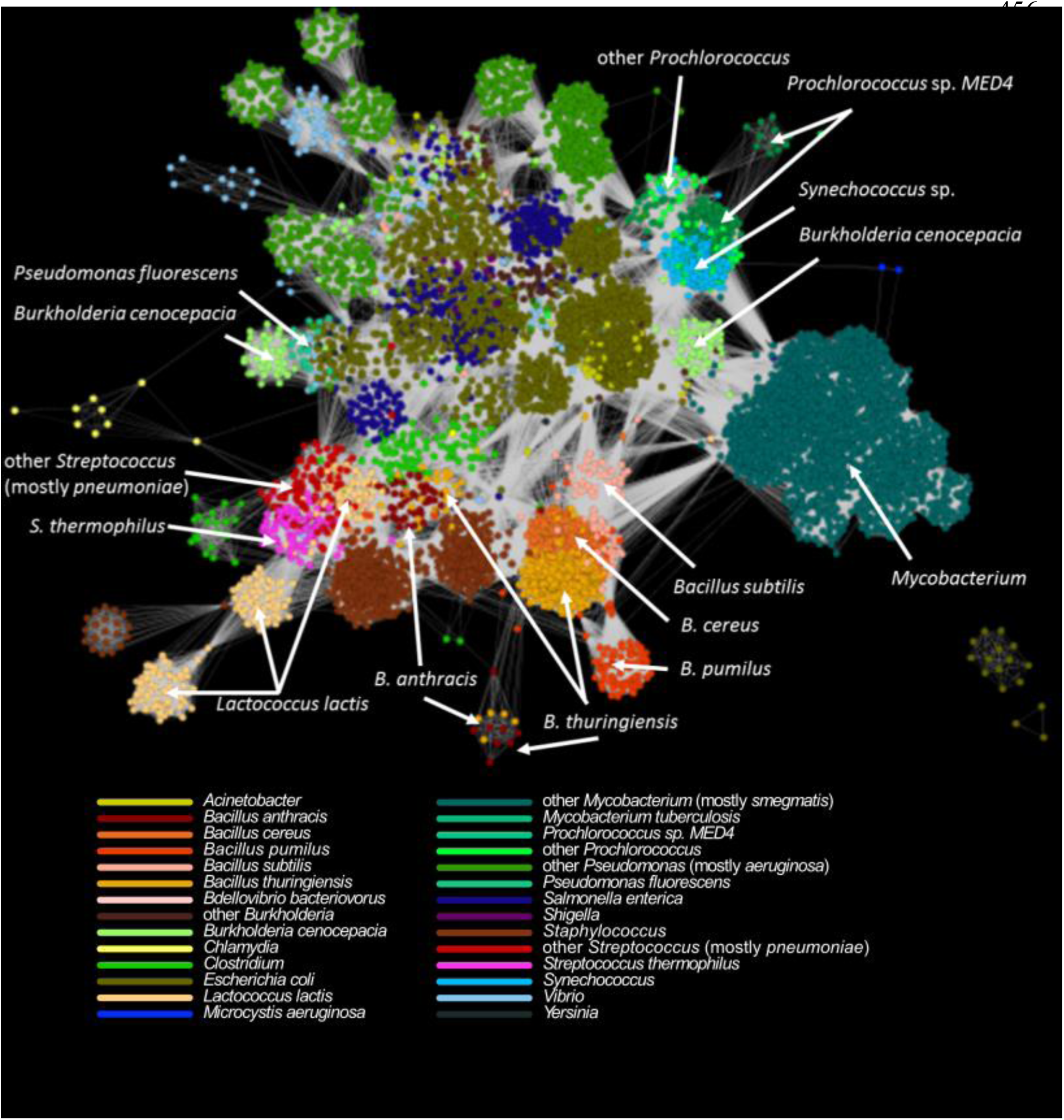
The gene network shown is identical to the network in Figures 1b and 2b, but with nodes recolored according to the host species, where annotation was available. Labels and arrows indicate specific cases highlighted in the main text.

In the case of *Bacillus* phages, genes are found in clusters corresponding to their annotated host species: *B. anthracis, B. subtilis*, *B. thuringiensis*, *B. pumilus*, or *B. cereus*. Further, overlap exists between *B. anthracis* and *B. thuringiensis*, closely related pathogens belonging to the *B. cereus* group (Priest et al. 2004). Host associations at the species level are also visible within the genera *Prochlorococcus* and *Streptococcus*.

Not all graphical clusters, however, correspond to a specific host species or strain. *Lactococcus lactis*, for instance, is frequently used in dairy starter cultures as *L. lactis* subsp. *lactis* and *L. lactis* subsp. *cremoris*, and phages have been well-sampled from both hosts (Deveau et al. 2006). Genes from these diverse phages occur across three clusters of phage genetic diversity in the gene network, with no clear associations with either host subspecies. Notably, these phages often are found to infect multiple strains of *L. lactis* (Mahony et al. 2013), and recombination between dairy phages may be frequent (Brüssow and Desiere 2001). Interestingly, one cluster of *Lactococcus-*associated genes shares many connections with a cluster of *Streptococcus thermophilus,* another common member of dairy fermentations.

The largest and most distinct cluster of phage genes corresponds to phages infecting *Mycobacterium smegmatis*, a non-pathogenic and more readily-cultured relative of *M. tuberculosis*. These phages have been heavily sampled compared to other hosts because of the SEA-PHAGES program, in which undergraduates isolate and sequence phage genomes (Jordan et al. 2014). Though phages of other species of *Mycobacterium* have not been thoroughly studied, genes from phages infecting *M. tuberculosis* are also present across MCL clusters found within this subgraph. This observation suggests it may be worthwhile, if technically difficult, to test more of the phages of *M. smegmatis* on *M. tuberculosis*, as has been previously suggested (Hatfull 2014).

Though not as well-sampled as phages of *Mycobacterium*, genes from phages infecting *Escherichia coli* and *Pseudomonas* species appear across the network, often more closely related to phages infecting other genera. Genes from phages of *Salmonella*, *Shigella*, *Acinetobacter*, and generically-identified *Enterobacteria* can all be found within clusters that are largely associated with *E. coli*. There is a distinct cluster of phage genes affiliated with *Pseduomonas fluorescens*, but other species-specific designations are not readily-observed. Iranzo *et al*. (2016) recently introduced a bipartite network connecting phage genes to phage genomes, which may provide further insight into how recombination events have structured phage host range.

## Quantifying and optimizing associations between network clusters and phage hosts

We next sought to quantify the reliability of these visible associations and to ask if subsets of genes could be used to predict a phage’s host. We estimated the degree of overlap between graphical clusters and host associations in each network by determining their mutual information (see Supplemental Methods). This metric suggested that clusters in the genome network may, in fact, be more closely associated with host annotations than clusters in the gene network (MI_genome_ = 2.18, MI_gene_ = 1.42). This effect likely arises, however, because each node in the genome network corresponds to exactly one host, and each MCL cluster in the genome network has, on average, only 4.36 members. In contrast, there are an average of 65.5 genes within each MCL cluster in the gene network, and each node within these clusters corresponds to at least 3 homologous genes from different phage genomes. More importantly, many genes are not directly linked to host specificity, and homologs represented by a single node in the gene network may come from phages that infect different hosts. Thus, graphical clusters built from the gene network will contain many genes with variable host associations, whereas those within the genome network are buffered from this noise. In the gene network, this variation reduces the mutual information between cluster membership and host. This effect would also imply that there exists a subset of genes within the gene network that would provide greater correspondence with host associations.

To address this hypothesis, we developed an evolutionary algorithm, *mimax*, to identify the subset of genes that maximizes the mutual information of MCL clusters and hosts. The *mimax* algorithm works as follows: in each iteration, an MCL cluster in the gene network is removed from a matrix of cluster-host associations at random. If doing so would result in removing a phage genome from the dataset, the deletion is rejected. If no genomes are lost, then the mutual information of the new matrix is calculated. If this value exceeds the value from the previous iteration, the deletion is retained, otherwise it is rejected. Because the *mimax* algorithm depends on removing uninformative clusters of genes, it should be more effective when there are more clusters from which to choose. When applied to the 135 clusters previously found in the gene network, *mimax* removed 47 clusters containing 1375 genes (~15% of the dataset), resulting in a modest improvement in mutual information but still falling short of the value observed in the genome network.

Three methods have been suggested to increase the granularity of MCL clusters (see https://micans.org): increasing the inflation factor, removing highly connected nodes before finding clusters, and introducing noise to the network. Initially, we chose an inflation factor of 4.1 to optimize the ICCC, a measure of within-cluster cohesion. The ICCC, though, is largely of interest when clusters represent naturally distinct sets of nodes, such as for taxonomic classification using genome-level networks. Here, we are more interested in subdividing genes into co-occurring subsets, and optimizing ICCC comes at the cost of sensitivity for the *mimax* algorithm. We tested each of the three methods described above (see Supplemental Methods and Figure S2), finding the best results, 1355 clusters, with an inflation factor of 15 and adding 5 random edges per node. Given this new set of clusters, we ran *mimax* 10 times and retained the resulting matrix with the highest mutual information. In each replicate, the mutual information between MCL membership and host associations converged to a higher value than found in the genome network (Figure S3). On average, *mimax* reduced the number of MCL clusters and associated genes within the gene network to 483.5 and 4070.6, respectively. These deletions suggest that over half of the genes in the gene network are uninformative with respect to host range.

Two questions emerge from maximizing the mutual information between graphical clusters and host associations: 1) Are the retained genes more closely associated with functions characteristic of phage-host interactions? and 2) can the resulting gene network be used as a tool for predicting the primary host of phages?

To address the first question, we annotated the complete and *mimax*-reduced sets of genes using RAST (Aziz et al. 2008). We then compared the frequency of common annotations of non-hypothetical proteins for each set of genes (see Table S1 and Figure S3). Phage baseplate, neck, replication, and DNA synthesis genes are over-represented following *mimax*, whereas phage packaging and regulatory genes are under-represented. Phage baseplate proteins directly affect virus adsorption to host receptors (Mahony and van Sinderen 2015), suggesting that gene function does affect *mimax* results.

The cluster-host correspondence in the *mimax*-reduced gene network offers a novel means to predict a phage’s host. Given a phage’s genome, we identified all genes that belong to the *mimax*-reduced set. We then recorded how often each potential host was associated with a homolog of one of these remaining genes (excluding a phage’s own contribution if already within the network). Finally, we chose the most frequent host affiliated with this subset of genes as the predicted host. (See the Supplemental Methods for additional details of the procedure.) When applied to all phage in the network, this approach predicted the host genus with 86% accuracy. If the full gene network is used in place of the *mimax*-reduced network, accuracy declines to 72%. This difference confirms that the *mimax* procedure reduces the gene network to a set of genes with stronger ties to phage host determination.

We deconstructed the host prediction accuracy (from *mimax*-improved predictions) for each host genus in order to account for uneven sampling of phages across hosts (Table 1). Doing so indicates that accuracy varied with host genus. Predictions for phages of *Mycobacterium* were nearly 100% accurate, and this reflects the large, unique space occupied by their genes in the network. In contrast, host predictions were less accurate for hosts with few representatives in the dataset, such as *Clostridium* and *Yersinia*. Accuracy also declined for well-sampled hosts, such as *Escherichia*, where phages have been sampled from closely-related genera (e.g. *Salmonella, Shigella,* and *Yersinia*). As has been seen for other host prediction methods (e.g. Villaroel et al. 2016), incorrect host predictions tended to predict that phage infect closely-related hosts (see Table 1). Improving the accuracy of predictions within these groups requires additional sampling and wet lab characterization of phages from across host genera. We should also be careful when assessing the quality of negative predictions. While phage host range can be exceptionally specific, many phages infect multiple genera (Hamdi et al. 2017, Jensen et al. 1998) or even across phyla (Malki et al. 2015), and additional lab work is required to confirm that putatively incorrect predictions are not, in fact, false negative results.

**Table 1:** Host accuracy varies with genus and sampling

| Host Genus | Accuracy | Top Mistake | Total Count |
| --- | --- | --- | --- |
| *Chlamydia* | 1 | N/A | 4 |
| *Lactococcus* | 1 | N/A | 36 |
| *Mycobacterium* | 0.991 | *Lactococcus* | 226 |
| *Bacillus* | 0.97 | *Chlamydia* | 66 |
| *Streptococcus* | 0.947 | *Bacillus* | 38 |
| *Escherichia* | 0.906 | *Salmonella* | 138 |
| *Prochlorococcus* | 0.905 | *Synechococcus* | 21 |
| *Staphylococcus* | 0.897 | *Bacillus* | 87 |
| *Pseudomonas* | 0.847 | *Escherichia* | 85 |
| *Burkholderia* | 0.833 | *Pseudomonas* | 30 |
| *Salmonella* | 0.804 | *Escherichia* | 56 |
| *Vibrio* | 0.686 | *Escherichia* | 51 |
| *Clostridium* | 0.667 | *Streptococcus* | 21 |
| *Acinetobacter* | 0.583 | *Escherichia* | 12 |
| *Shigella* | 0.273 | *Escherichia* | 11 |
| *Yersinia* | 0.273 | *Escherichia* | 11 |
| *Anabaena* | 0 | *Escherichia* | 1 |
| *Microcystis* | 0 | *Escherichia* | 1 |
| *Chlamydophila* | 0 | *Chlamydia* | 1 |
| *Synechococcus* | 0 | *Prochlorococcus* | 15 |
| *Bdellovibrio* | 0 | *Escherichia* | 2 |

We next tested this approach with a set of novel phages not included in the original gene network. Over 1000 new phage genomes have been published since we built our original network. We chose 500 phage genomes at random from this new set. Of these, 185 were annotated as infecting hosts already included in our network. The genes in these phages were assigned to the *mimax*-reduced set of MCL clusters identified previously. While 52 of these phages shared no genes in the *mimax* set with any phages in our original dataset, for the remaining 133 phage, our procedure predicted the host genus 67.7% of the time (see Table S2). Moreover, accuracy remained high for well-sampled hosts, such as *Mycobacterium* and *Escherichia*, but was low for others, such as *Bacillus*, that we could previously predict with over 90% accuracy. This discrepancy suggests that a gene network approach to host prediction should be updated regularly to account for the frequent addition of new virus genomes to repositories.

## Conclusion

In this work, we have shown that gene-level networks provide both a high-resolution view of viral genetic diversity and a means to connect specific groups of genes to broad patterns in viral ecology. When applied to virus host range, phage gene clusters correlated with a phage’s annotated host, and proximity of clusters in the network reflected the evolutionary relatedness of these hosts. Using an evolutionary algorithm, *mimax*, we were then able to identify specific groups of genes with the strongest correlation to virus host range. The *mimax*-reduced dataset was enriched for genes known to affect host recognition, and the enhanced network offers one of the most accurate means of host prediction to date.

This approach should be extensible to aspects of viral ecology beyond host range, including isolation source (e.g. freshwater, marine, soil, leaf, gut, hospital, etc.) and abiotic or biotic factors that vary across locations (e.g. temperature, pH, O2, nutrient concentrations, and available host diversity). Moreover, phage have a direct impact on the growth of their host bacteria, and knowing a phage’s ecological and evolutionary history is critical to understanding how that phage affects an ecosystem. Gene network analysis should facilitate new discoveries in any environment, be it a dairy vat, a freshwater lake, or the human gut.

## Acknowledgments

We are grateful to past and present members of the Putonti lab for thoughtful feedback on this work.

